# A Bayesian network approach for modeling mixed features in TCGA ovarian cancer data

**DOI:** 10.1101/033332

**Authors:** Qingyang Zhang, Ji-Ping Wang, Northwestern PSOC members

**Affiliations:** Q. Zhang is with Department of Statistics, Northwestern University, Evanston, IL 60208,.; J.-P. Wang is with Department of Statistics, Northwestern University, Evanston, IL 60208,.

## Abstract

We propose an integrative framework to select important genetic and epigenetic features related to ovarian cancer and to quantify the causal relationships among these features using a logistic Bayesian network model based on *The Cancer Genome Atlas* data. The constructed Bayesian network has identified four gene clusters of distinct cellular functions, 13 driver genes, as well as some new biological pathways which may shed new light into the molecular mechanisms of ovarian cancer.

## I. Introduction

While the molecular mechanism of ovarian cancer remains unclear, studies have suggested that many different factors may contribute to this disease, among which there are tens of well-known oncogenes and tumor suppressors including *TP53, PIK3C, PTEN, BRCA1* and *BRCA2* [1,2]. However, the analysis based on individual genes often fails to provide even moderate prediction accuracy of the cancer status. Thus a systems biology approach that combines multiple genetic and epigenetic profiles for an integrative analysis provides a new direction to study the regulatory network associated with ovarian cancer. Here we describe an integrative framework that presents two innovations: (1) a novel stepwise correlation-based selector (SCBS) for important pathway-relevant features; and (2) a Bayesian network (BN) modeling for a mixture of continuous and categorical variables for casual inference. This approach provides a way to mine the massive cancer data for important genetic and epigenetic features directly or indirectly associated with cancer phenotype, leading to discoveries of pathways underlying the molecular mechanism of cancers.

## II. Results

### 1. Predicted Bayesian network

We consider four important data types in our analysis in the TCGA ovarian cancer data ([1], Table 1). First, 68 potential oncogenes and tumor suppressors (identified from both literature [2] and TCGA data) were used as seed genes. An additional 271 nodes out of more than 50,000 candidate features were selected by the SCBS (Methods and [3]), including expression level of 177 genes, 82 copy number variation sites, 11 methylation sites and one somatic mutation site at gene *TP53*. The predicted BN (Fig. 1) contains many well-known biological pathways. To name a few, the edge from *CDKN2A* to *CCNE1* is a known genegene regulation in the RB signaling pathway [1]. The edge from *TPX2* to *AURKA* confirms that *TPX2* can activate *AURKA* by inducing autophosphorylation [4].

**Table 1:**
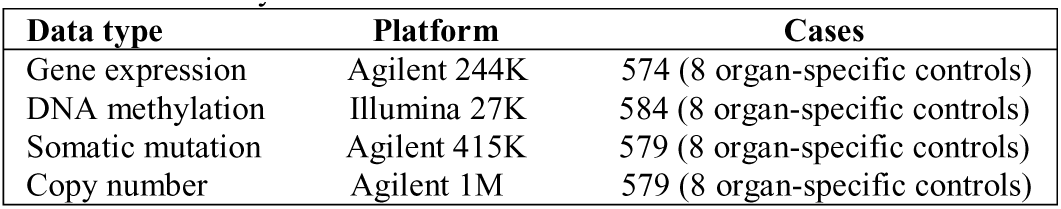
Summary of TCGA ovarian cancer data

| Data type | Platform | Cases |
| --- | --- | --- |
| Gene expression | Agilent 244K | 574 (8 organ-specific controls) |
| DNA methylation | Illumina 27K | 584 (8 organ-specific controls) |
| Somatic mutation | Agilent 415K | 579 (8 organ-specific controls) |
| Copy number | Agilent 1M | 579 (8 organ-specific controls) |

**Figure 1:**
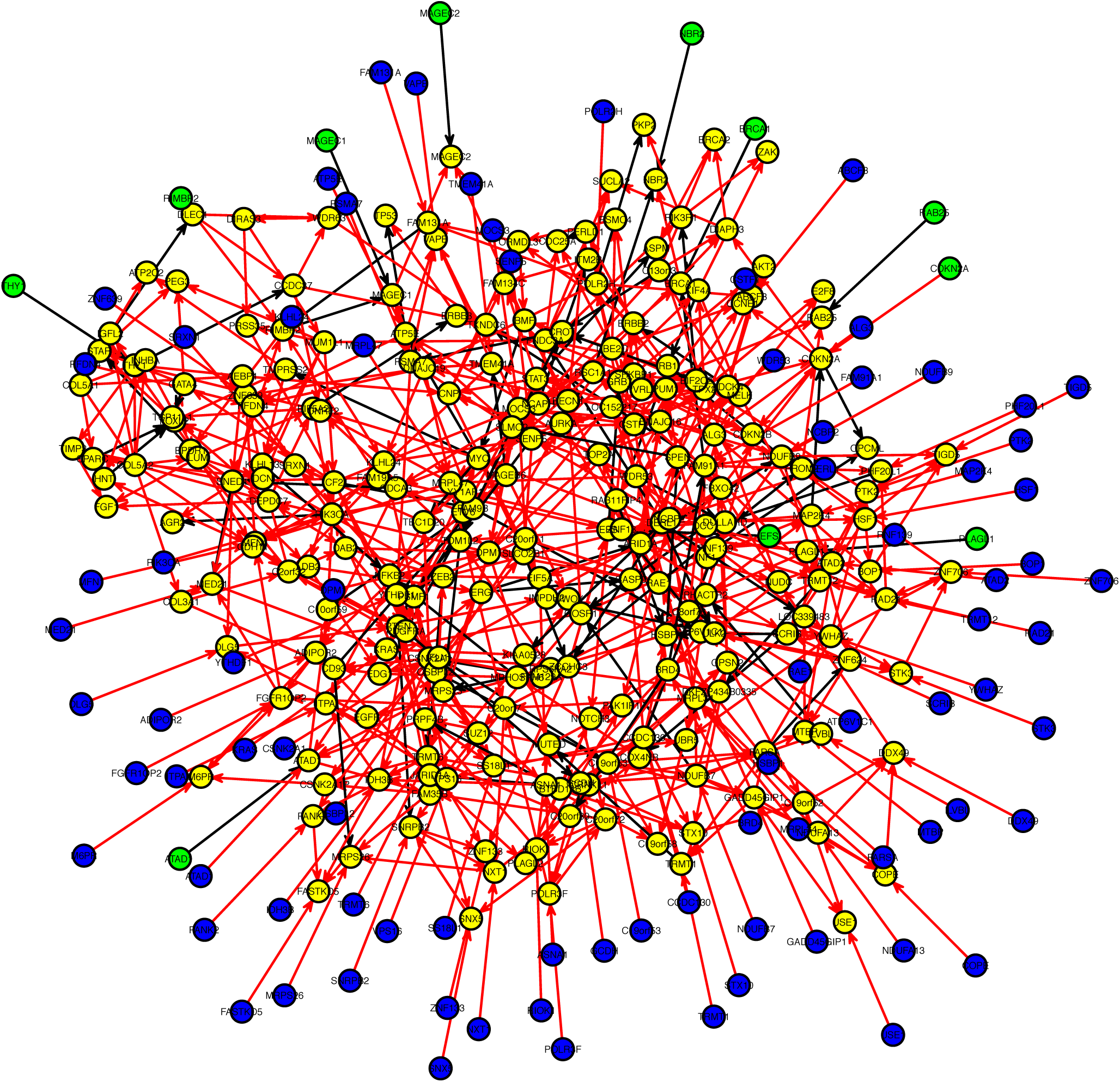
Predicted graph by logistic BN model with 339 nodes including expression levels of 245 genes (yellow), copy number at 82 sites (blue), methylation at 11 sites (green) and 1 somatic mutation at *TP53*. Direction of edges means the downstream feature is regulated by the upstream one. Red edges represent activation and black edges represent suppression.

### 2. Hub genes and sample classification

We identified 13 nodes with significantly larger out-degrees in the network: *ARID1A, C19orf53, CSNK2A1, DERL1, TRMT6, COL5A2, TCF21, LUM, TPX2, UBE2C*, DPM1, *NDUFB7* and *NDUFB9*. We show (in Fig. 2) that the 13 hub genes can clearly distinguish the cancer samples from the normal samples by a multi-dimensional scaling (MDS) plot based on the correlation dissimilarity metric (comparable clustering effect was observed based on the entire set of 245 genes). This suggests that the thirteen hub genes may present the major difference between the cancer and normal samples. The early-stage and high-grade tumor samples however are not well distinguished.

**Figure 2:**
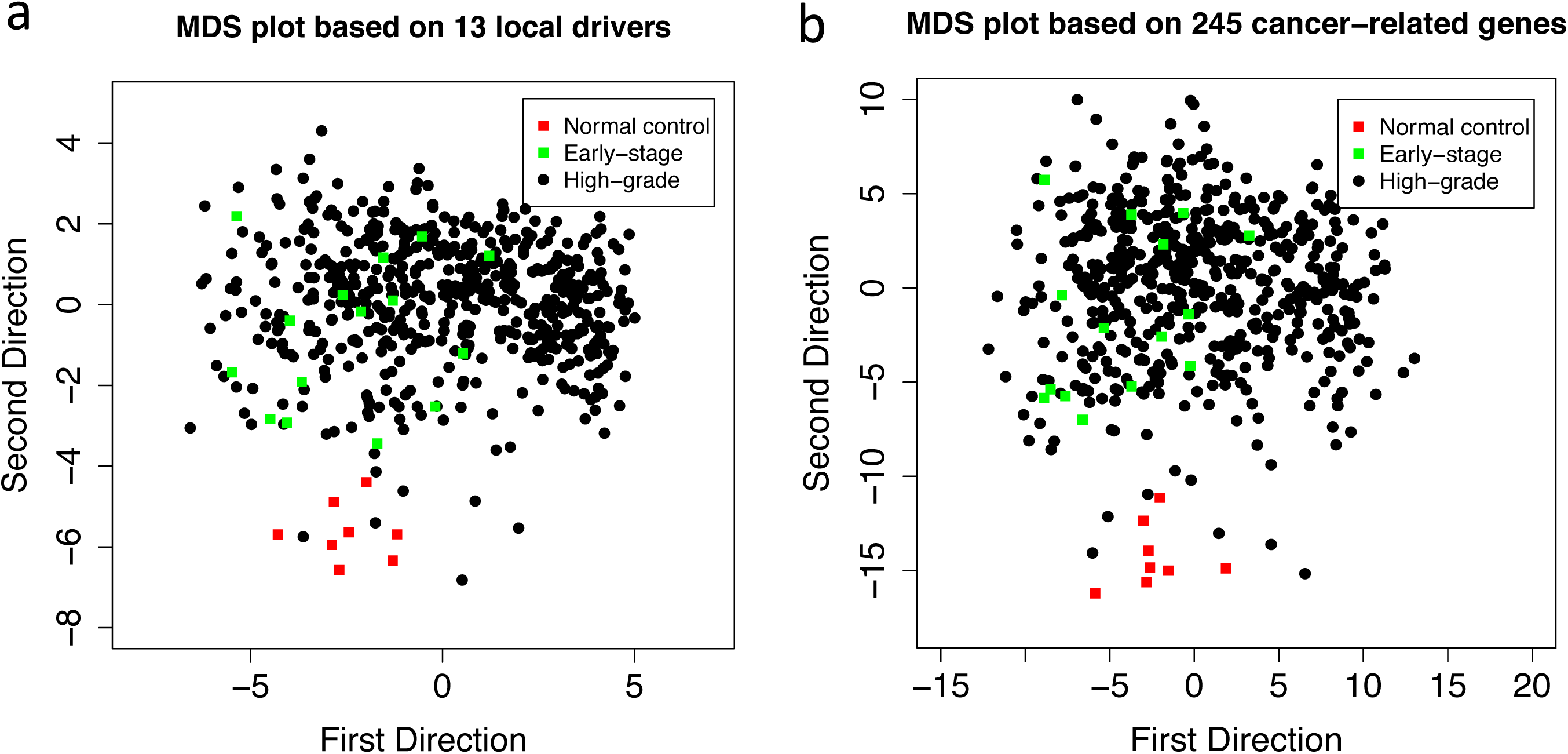
MDS plots for sample classification. (a) based on 13 hub genes; (b) based on all 245 genes in the predicted network. Each dot represent on sample: red (cancer-free), green (early-stage cancer) and black (high-grade cancer).

### 3. Major Gene clusters

The 245 genes fell into four major clusters corresponding to distinct functions by k-means clustering method [3]. Cluster 1 (red, Figure 3a) contained 18 genes, mainly related to cell division, mitosis, spindle formation etc. Cluster 2 (green, Figure 3a) contained 23 genes, most of which are functionally related to growth factor, cell shape, cell motility, tumor invasion etc. Cluster 3 (black, Figure 3a) contained 20 genes, mostly related to mitochondrial system, membrane process etc. Cluster 4 (blue, Figure 3a) was the largest and most complicated cluster harboring 184 genes. This large cluster communicates between the other three clusters, which were nearly independent from each other (Figure 3b). These findings could be implicative of some important molecular pathways, which may or may not have been identified, that drive the development of ovarian cancer.

**Figure 3:**
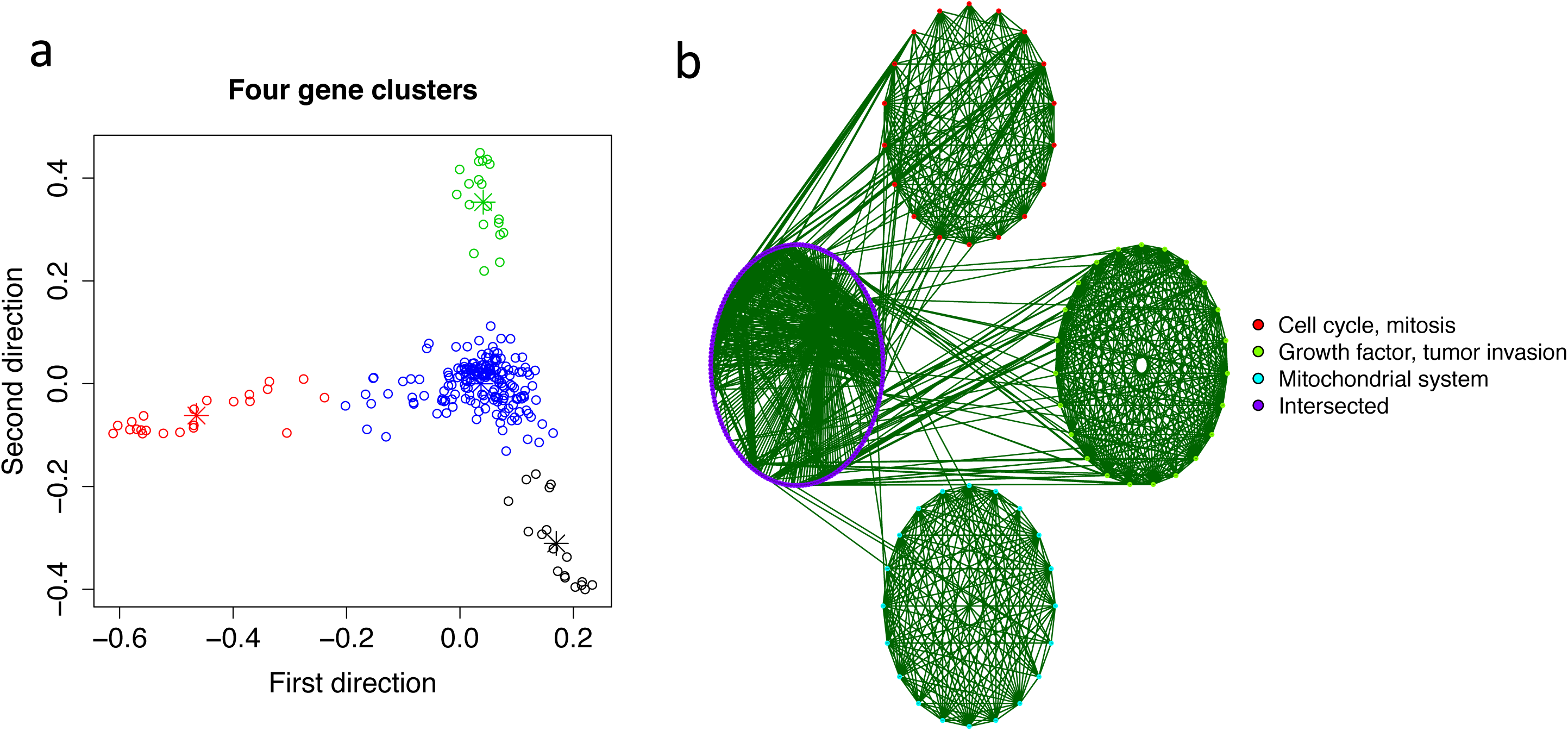
(a) MDS plot based on correlation dissimilarity metric between 245 genes (each circle represents one gene). Genes falling into four clusters (by k-means clustering method where k = 4) are indicated by different colors; (b) Correlation plot of the four clusters, the connection between a pair of genes represents a significant correlation.

## III. Methods

### 1. Bayesian network with logit link function

The Bayesian network is one way to model the causal relationships among a set of random variables (nodes) via a directed acyclic graph ([5]). In general, the joint likelihood of nodes *X*_1_, *X*_2_,…,*X_p_* can be expressed as follows:

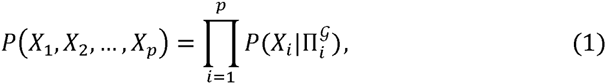

where 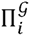 stands for the parental nodes of node *i* in the graph. The proposed logistic BN model ([3]) incorporates both continuous and discrete variables (require discretization of the continuous variables). For simplicity, we illustrate the model under binomial case:

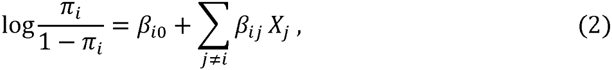

where *X_i_* takes value 0 or 1 with respective probabilities {1 – *π_i_*, *π_i_* }. Here we transform the network structure to the coefficient matrix {*β_ij_* }, where = 0 means *X_j_* ⇸ *X_i_* (no causal effect form node *j* to *i*) and otherwise *X_j_* → *X_i_* (with causal effect). The details of the model fitting can be found in Zhang et al [3].

### 2. Stepwise correlation-based feature selector

One important and necessary step before BN learning is feature selection. The prevailing method is based on independent test, which can falsely selects many irrelevant features as some features could be causal to other features while having no direct association with the cancer phenotypes. With the SCBS procedure, we start with detection of features significantly correlated with the cancer and then progressively select subsequent features that correlate with the selected features. Suppose we aim to select *p* variables (out of *S* candidates, *S* ≫ *p*) as the nodes in BN based on *N* random samples. SCBS can be implemented as follows:

Step 1: Calculate the correlation coefficients between the current node *X_i_* and all the other nodes. Keep *k* most correlated nodes with *X_i_* for further filtering.
Step 2: Calculate the *p*-value of correlation coefficient for each of the *k* nodes from step 1, Select the node if the *p*-value is significant under Benjamini-Hochberg procedure with FDR ≤ 0.05.
Step 3: Repeat step 1 and 2 until *p* nodes are selected.

For the choice of *k*, we suggest *k* = 4, 5 or 6 in practice since smaller *k* tends to miss weakly connected nodes and larger *k* tends to catch more false positives. See a simulation study in [3] for detail.

## Notes

Research supported by NCI grant #U54CA143869

## REFERENCES

[1] The Cancer Genome Atlas Research Network, “Integrated genome analyses of ovarian carcinoma”, Nature, vol. 474, pp. 609–615

[2] R. Bast, B. Hennessy and G. Mills, “The biology of ovarian cancer: new opportunities for translation”, Nature Cancer Review, vol. 9, pp. 415–428, 2009

[3] Q. Zhang, J.E. Burdette and J.-P. Wang, “Integrative network analysis of TCGA data for ovarian cancer”, BMC Systems Biology, vol. 8, No. 1338, pp. 1–18, 2014

[4] T. Kufer, H. Sillje, R. Korner, O. Gruss, P. Meraldi and E. Nigg, “Human TPX2 is required for targeting Aurora-A Kinase to the spindle”, Journal of Cell Biology, vol. 158, No. 4, pp. 617–623, 2002

[5] P. Spirtes, C. Glymour and R. Scheine, Computation, Causation and Discovery. Cambridge: Springer, 1993

